# A simple method for constructing magnetic *Escherichia coli*

**DOI:** 10.1101/010249

**Authors:** Mengyi Sun, Yong-jun Lu

**Affiliations:** School of Life Sciences and Biomedical Center, Sun Yat-sen University, No. 135, Xingang Xi Road, Guangzhou, 510275, P. R. China

**Keywords:** Ammonium ferric citrate, *E. coli*; FtnA, Magnetization

## Abstract

Magnetic force can serve as an ideal way to control the spatial behavior of microorganisms, because of its flexibility and penetrability. By incubation with the biocompatible compound, ammonium ferric citrate, as an iron source, we magnetized *Escherichia coli*, the most programmable chassis in synthetic biology. To enhance the magnetization efficiency, the ferritin protein, FtnA, from *E. coli* was cloned and overexpressed in strain BL21(DE3). The magnetization effect was observed within 30 min after harvest of bacteria, and the concentration of ammonium ferric acid used could be as low as 0.5 mM. Using different shapes of magnetic fields, different patterns could be generated easily. Our method may set up the foundation for a rational design of spatial structure of cell communities, which is important for their actual application.

## Introduction

Controlling the movement and location of cells is one of the most important topics in synthetic biology. Pattern formation, synthetic consortia and the novel cell array sensor heavily rely on the precise spatial arrangement of cells (Melamed et al. 2012; Brenner et al. 2008). The traditional method relied on modifying the chemotaxis mechanism of cells, which was both slow and irreversible (Goldberg et al. 2009). Recently, rapid control of cells has been achieved by introducing light controllable proteins into organisms, hence the name optogenetics (Herrou and Crosson 2011). However, the heat effect of light prevented its use in prolonged control. Furthermore, the low penetrability of light hindered its application. Compared with chemical and light control, magnetic force is safe, fast, reversible, highly penetrative and easy to design (Pankhurst et al. 2003), and thus would be an ideal method for controlling cells if the cells could be magnetized. Therefore, several methods to magnetize cells have been developed, including methods that rely on artificial synthesis of magnetic nanoparticles (Cho et al. 2012), and transferring the cassette controlling magnetosome synthesis in *Magnetospirillum gryphiswaldense* into the related species, *Rhodospirillum rubrum* (Kolinko et al. 2014). However, these methods are both laborious and hard to generalize. In 2011, Silver’s group (Nishida and Silver 2012) reported a simple method for magnetizing budding yeast, by incubation with ferric citrate and introduction of a ferritin protein.

So far, no simple method for magnetizing the most programmable chassis organism in synthetic biology has been reported. Here, we extended Silver’s method to *Escherichia coli*. After incubation with ammonium ferric citrate, we showed the magnetization by the formation of defined patterns under strong magnets. By cloning and overexpressing FtnA, a ferritin protein that helps store iron in *E. coli* (Bitoun et al. 2008), we showed that the magnetization effect could be amplified even under a low concentration of Fe^3+^. Using different shapes of magnets, we also showed the flexibility of patterns formed by bacteria. Our method provides a versatile tool for spatial control of *E. coli*, which may aid the development of synthetic biology.

## Methods

### Strains, culture medium and conditions

The *E. coli* strains used for cloning FtnA, plasmid construction and the attraction test were K12, DH5α and BL21(DE3), respectively. The medium used was LB. Ampicillin was added when necessary. Except for induction, the bacteria were cultured at 37°C with agitation at 220 rpm.

### Plasmid construction

The *ftnA* gene was amplified from strain K12 and cloned into plasmid pUC18. A mutation was introduced to eliminate the NdeI restriction site in the original *ftnA* sequence using the QuickChange kit (Stratagene). Then, the coding sequence was ligated into the pET-22b+ vector using the NdeI and BamHI restriction sites (Fig. 1).

**Fig. 1.**
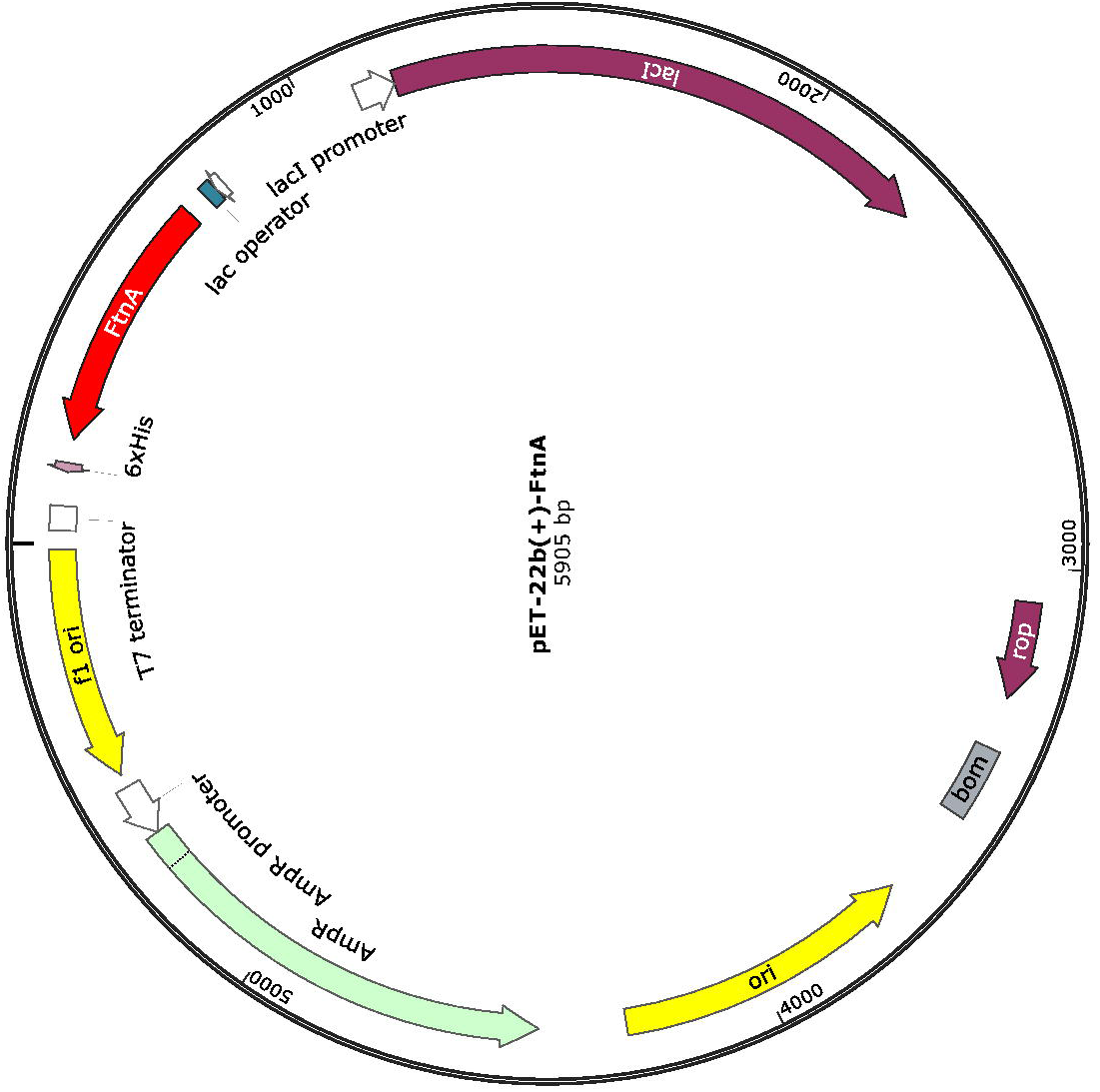
The map of plasmid pET-22b+-FtnA, which was constructed by inserting the FtnA coding sequence immediately after the ribosomal binding site

### Growth test

Experiments were performed at 37°C with agitation at 220 rpm. OD_600_ values were measured at several time points, and converted into dry weight. The experiments were performed in triplicate.

### Magnetic cells preparation

For attraction tests without plasmid induction, bacteria were cultured in LB medium inoculated with 20 mM Fe^3+^ derived from ammonium ferric citrate. Cells were harvested after 24 h and washed several times, then pelleted by centrifugation and resuspended in double-distilled H_2_O. For induction experiments, cells were exposed to 20 mM Fe^3+^ and induced with 1 mM isopropyl β-D-1-thiogalactopyranoside (IPTG) at 25°C for 24 h, and then harvested as described above.

### Attraction test

Three milliliters of cell suspension were added to 3.5-cm dishes. For characterizing the magnetization effect, a circular Rubidium iron boron magnet (2-cm in diameter) was put underneath each dish, representing a magnetic field of 1.2 T near the magnet edge. For generating different patterns, different shapes of magnets were used, accordingly. Photographs were taken at appropriate time points.

## Results and Discussion

### Effect of ammonium ferric citrate on ***E. coli* growth**

To obtain magnetic *E. coli* as much as possible, we used ammonium ferric citrate, since citrate has been reported as an effective microorganism siderophore for iron absorption (Guerinot et al. 1990). We compared the growth of *E. coli* under high iron concentration (20 mM ammonium ferric citrate added from a fresh stock of 500 mM) and normal conditions by measuring the OD_600_ at several time points after inoculation. As shown in Fig. 2, no significant difference was observed between the control group and the experimental group, demonstrating that ammonium ferric citrate is not toxic to *E. coli* cells. Hence, ammonium ferric citrate could serve as the iron source without decreasing the amount of *E. coli* harvested. This enabled us to easily maintain a population that is much larger than any other chassis organism for a synthetic cell-cell community (Youk and Lim 2014).

**Fig. 2.**
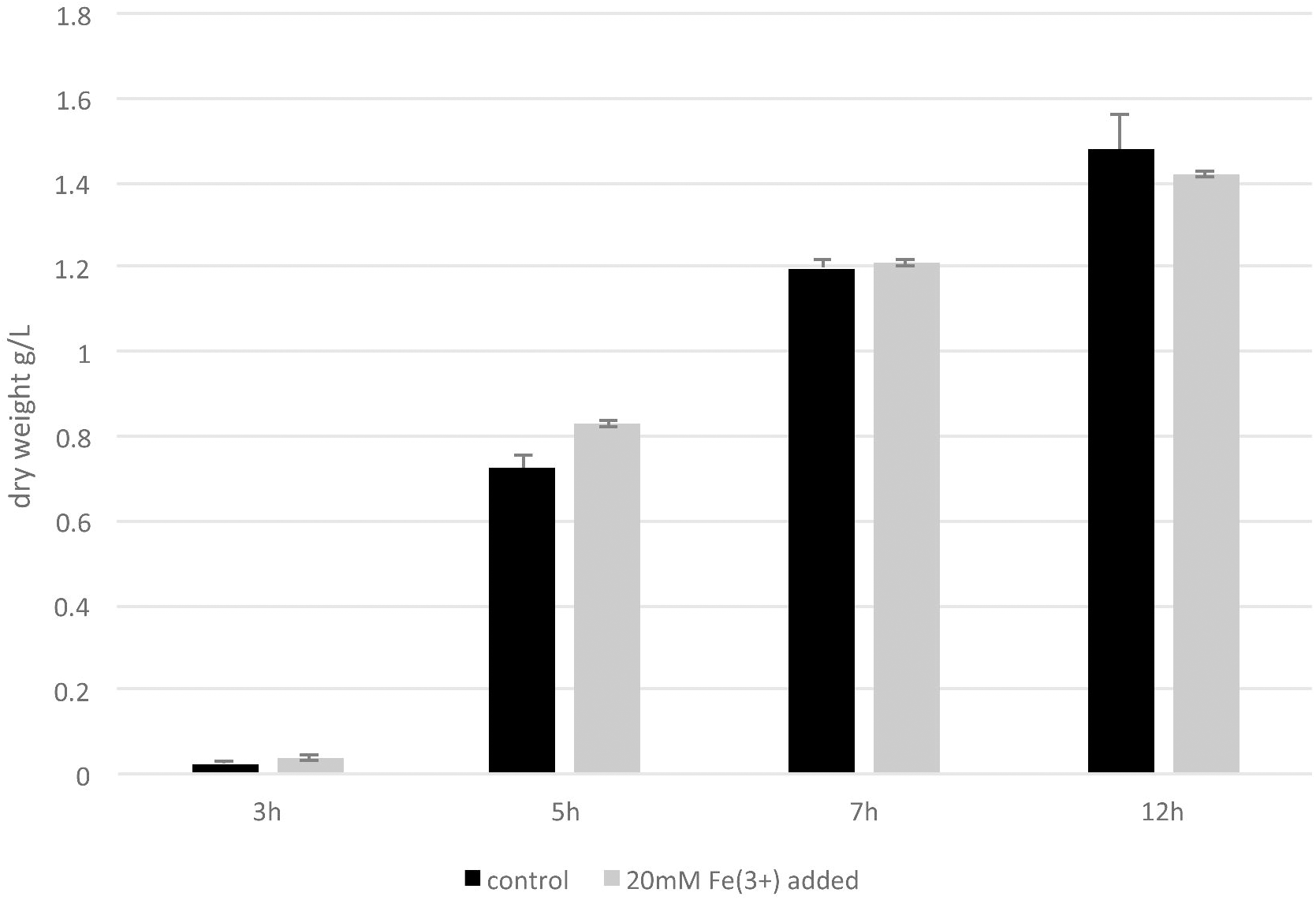
Growth test of *E. coli* inoculated with ammonium ferric acid. The OD_600_ values were measured at 3, 5, 7 and 12 h. Experiments were performed in triplicate

### Magnetization of *E. coli* by incubation with ammonium ferric citrate

Next, we tested whether *E. coli* could be magnetized by incubation with ammonium ferric citrate. Cells were harvested 24 h after inoculation. As shown in Fig. 3a, cells harvested from cultures containing 20 mM Fe^3+^ formed a white circle where the magnetic force was strongest, near the edge of the magnet. No noticeable cluster of *E. coli* was formed in the control group. This result demonstrated that incubation with ammonium ferric citrate indeed magnetized *E. coli*. The short time and ease of preparation of the magnetic *E. coli* enabled a massive experiment to be performed in a short time.

**Fig. 3.**
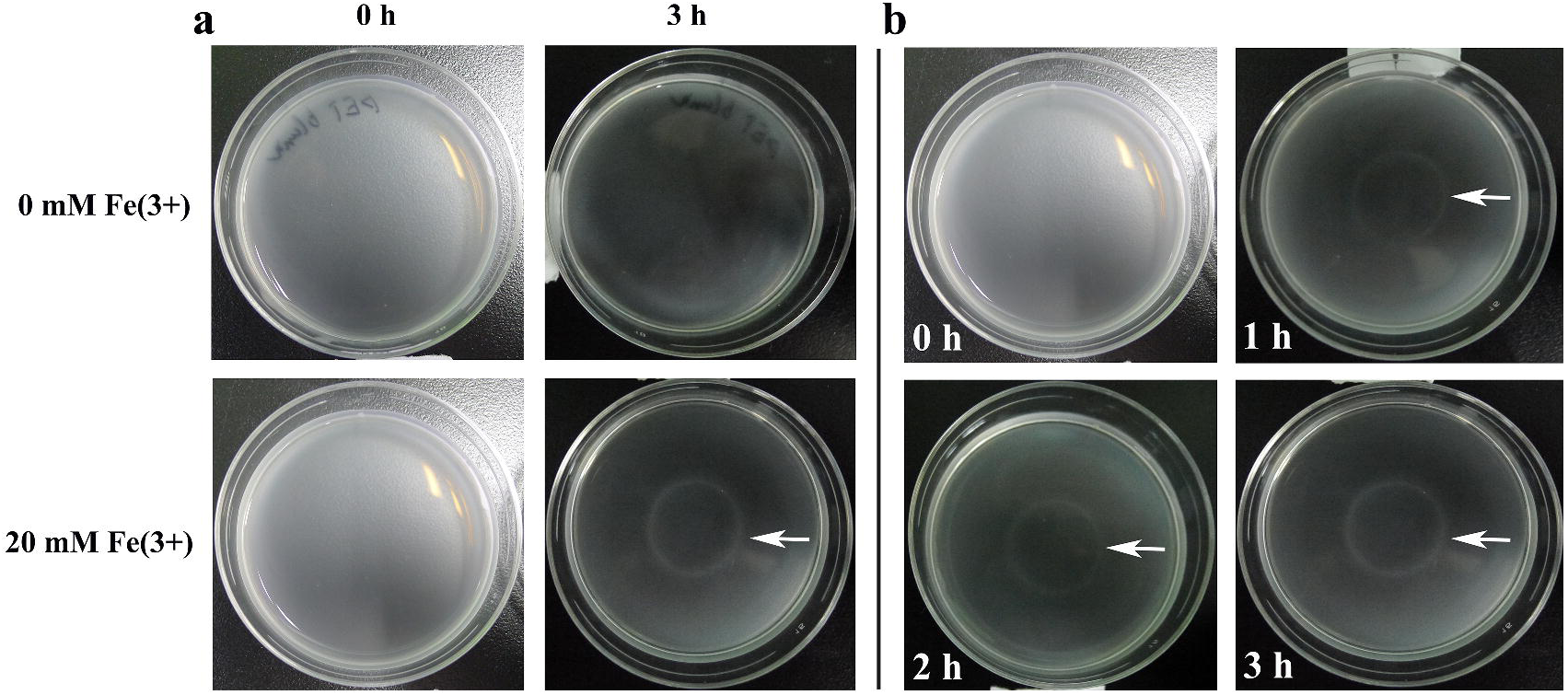
The test of the magnetization effect of inoculation with ammonium ferric acid by attraction ring formation. Bacteria resuspended in water were added to 3.5-cm blank dishes. A circular magnet was placed underneath each dish to attract magnetized bacteria. **(a)** The attraction ring was formed only in the group inoculated with ammonium ferric citrate. **(b)** A time course of the attraction ring formation

### The time course of the attraction phenomena

As shown in Fig. 3b, the attraction ring appeared 1 h after putting the magnet under the culture plate, and the pattern was stable after 2 h. Note that the pattern was formed in a liquid environment, which could not be accomplished by the traditional method with the chemotaxis mechanism, because the chemicals would quickly become uniform by diffusion. The *E. coli* could respond to the magnetic fields so quickly, because of the high efficiency of iron absorption of bacteria (Andrews et al. 2003). It is also well known that *E. coli* can express ferritin, a ubiquitous iron storage protein that is conserved from bacterium to human. This protein becomes paramagnetic when iron is incorporated (Bauminger and Nowik 1989). Note that in our experiment, the time required to form the attraction ring was slightly longer than that in Silver’s experiment using yeast, probably because *E. coli* cells are much lighter than yeast cells (Nishida and Silver 2012).

### Improving the magnetization effect by introducing FtnA

To achieve a higher magnetization efficiency of *E. coli*, we cloned and overexpressed FtnA, a ferritin protein, in *E. coli* to enhance the efficiency of ferrite incorporation. This protein also increases the iron stores, which could be used during iron deficiency (Abdul-Tehrani et al. 1999). As shown in Fig. 4, the attraction ring was formed within 30 min in the experimental group, while in the control group no observable ring was formed in such a short time. Note that the concentration of Fe^3+^ used could be as low as 0.5 mM. Thus, the introduction of FtnA significantly increased the magnetization efficiency. This result also revealed a remarkable advantage of our system: the concentration of the iron source used could be as low as 0.5 mM, which was much lower than that required for magnetizing yeast (Nishida and Silver 2012).

**Fig. 4.**
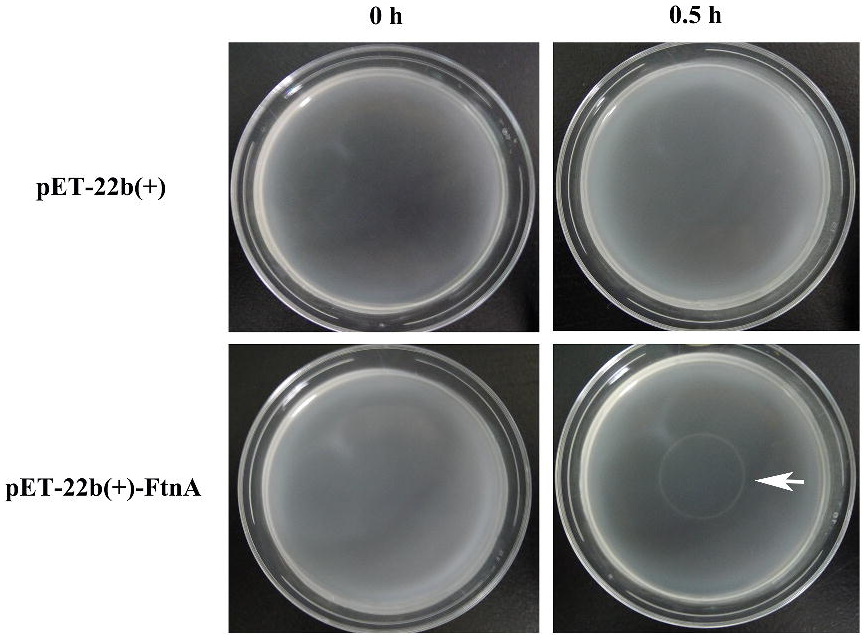
Overexpressing FtnA improves the magnetization effect

### The flexibility of the pattern formed by magnets of different shapes

To show the flexibility of the cell pattern formation, we used different shapes of magnets to induce different pattern shapes. As shown in Fig. 5, a rectangular shape and small double circles were formed under the attraction of rectangular magnets and small circular magnets, respectively. Interestingly, our method generated accurate pattern edges comparable to the synthetic edge detection circuit in *E. coli* (Tabor et al. 2009), without complex genetic design or mathematical modeling.

**Fig. 5.**
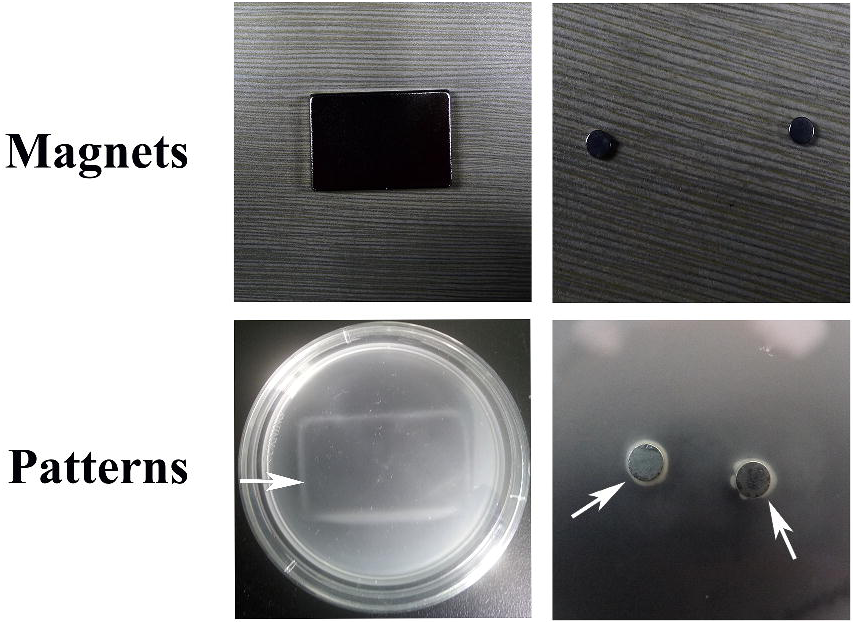
Different attraction rings with a sharp edge under the action of different magnet shapes

## Conclusion

Here, we established a simple method for producing magnetized *E. coli* by incubation with ammonium ferric citrate. The attraction ring pattern could be formed within 1 h. Overexpression of FtnA in *E. coli* significantly improved the magnetization effect. Furthermore, different patterns could easily be generated by different shapes of magnets, demonstrating the flexibility of our method. This method sheds light on using the magnetic force to control the most programmable microorganism, *E. coli*, in synthetic biology.

## Acknowledgments

We thank Dr. He Lei for his assistance with graphing, and members of XiongLei He’s lab for valuable suggestions and discussion. This work was supported by NSFC (No. J1310025).

The authors declare no conflict of interest.

